# Rare protein coding variants implicate genes involved in risk of suicide death

**DOI:** 10.1101/2020.01.10.902304

**Authors:** Emily DiBlasi, Andrey A. Shabalin, Eric T. Monson, Brooks R. Keeshin, Amanda V. Bakian, Anne V. Kirby, Elliott Ferris, Danli Chen, Nancy William, Eoin Gaj, Michael Klein, Leslie Jerominski, W. Brandon Callor, Erik Christensen, Douglas Gray, Ken R. Smith, Alison Fraser, Zhe Yu, PsychChip Investigators of the Psychiatric Genomics Consortium, Nicola J. Camp, Eli A. Stahl, Qingqin S. Li, Anna R. Docherty, Hilary Coon

**Affiliations:** Department of Psychiatry, University of Utah School of Medicine, Salt Lake City, UT, USA; Department of Pediatrics, University of Utah, Salt Lake City, UT, USA; Department of Occupational & Recreational Therapies, University of Utah, Salt Lake City, UT, USA; Department of Neurobiology & Anatomy, University of Utah School of Medicine, Salt Lake City, UT, USA; Health Sciences Center Core Research Facility, University of Utah, Salt Lake City, UT, USA; Utah State Office of the Medical Examiner, Utah Department of Health, Salt Lake City, UT; Pedigree & Population Resource, Huntsman Cancer Institute, University of Utah, Salt Lake City, UT, USA; Department of Internal Medicine, University of Utah School of Medicine, Salt Lake City, UT, USA; Pamela Sklar Division of Psychiatric Genomics, Icahn School of Medicine at Mount Sinai, New York, NY; Medical and Population Genetics, Broad Institute, Cambridge, MA; Research Information Technology, Janssen Research & Development LLC, Pennington, NJ; Virginia Institute for Psychiatric & Behavioral Genetics, Virginia Commonwealth School of Medicine, VA, USA

**Author notes:** Corresponding author; address for correspondence: Emily DiBlasi, University of Utah Department of Psychiatry, 383 Colorow Dr, University of Utah, Salt Lake City, Utah 84108.

## Abstract

Suicide death is a worldwide health crisis, claiming close to 800,000 lives per year. Recent evidence suggests that prediction and prevention challenges may be aided by discoveries of genetic risk factors. Here we focus on the role of rare (MAF <1%), putatively functional single nucleotide polymorphisms (SNPs) in suicide death using the large genetic resources available in the Utah Suicide Genetic Risk Study (USGRS). We conducted a single-variant association analysis of 30,377 rare putatively functional SNPs present on the PsychArray genotyping array in 2,672 USGRS suicides of non-Finnish European (NFE) ancestry and 51,583 publicly available NFE controls from gnomAD, with additional follow-up analyses using an independent control sample of 21,324 NFE controls from the Psychiatric Genomics Consortium. SNPs underwent rigorous quality control, and among SNPs meeting significance thresholds, we considered only those that were validated in sequence data. We identified five novel, high-impact, rare SNPs with significant associations with suicide death (*SNAPC1*, rs75418419; *TNKS1BP1*, rs143883793; *ADGRF5*, rs149197213; *PER1*, rs145053802; and *ESS2*, rs62223875). Both *PER1* and *SNAPC1* have other supporting gene-level evidence of suicide risk, and an association with bipolar disorder has been reported for *PER1* and with schizophrenia for *PER1, TNKS1BP1*, and *ESS2*. Three genes (*PER1, TNKS1BP1*, and *ADGRF5*), with additional genes implicated by GWAS studies on suicidal behavior, showed significant enrichment in immune system, homeostatic and signal transduction processes. Pain, depression, and accidental trauma were the most prevalent phenotypes in electronic medical record data for the categories assessed. These findings suggest an important role for rare variants in suicide risk and provide new insights into the genetic architecture of suicide death. Furthermore, we demonstrate the added utility of careful assessment of genotyping arrays in rare variant discovery.

## INTRODUCTION

Nearly 800,000 preventable suicide deaths occur each year.^1^ While environmental factors are undeniably important, evidence suggests that genetic factors play a major role in suicide death, with estimated heritability from meta-analyses of close to 50%.^2,3^ Identification of suicide genetic risk factors is critical to understanding the biological basis of suicide risk and improving prevention. However, to date only a fraction of the genetic variation influencing suicide risk has been accounted for with most studies focusing on suicidal behaviors (e.g. ideation or attempt) as opposed to suicide death.^4^

Most genetic studies of suicide focus on common genetic risk variants (variants carried by more than 1% of a given population), despite the high likelihood that rare variants (minor allele frequency; MAF <1%) also significantly contribute to suicide risk.^5,6^ Discoveries of rare genetic risk variants for other complex psychiatric and non-psychiatric medical conditions (e.g., lipid disorders^7^, Alzheimer’s disease^8^, schizophrenia^9^, obesity^10,11^) have identified functional gene pathways and now explain a sizeable proportion of the genetic risk for these disorders. Functional variants that alter gene function or expression tend to be rare in populations because natural selection acts to reduce their prevalence.^13^ Identification of risk genes and pathways affected by these rare but high-impact variants is critical to the development of targeted interventions.

A primary reason why most research focuses on common variants is that detection and reliable estimates of frequency of rare variants in both case and control groups requires very large genetic studies,^14,15^ often including many tens of thousands of carefully chosen samples to achieve statistical power.^16^ Whole genome sequencing (WGS) and whole exome sequencing (WES) studies are becoming more common, yet costs involved are considerable and these studies often have more modest sample sizes compared with genome-wide association studies (GWAS) using cheaper genotyping arrays. Large genetic datasets from genotyping arrays are an overlooked source of important rare variation. Although most of the content on genotyping arrays is focused on common variants, carefully chosen, medically relevant, rare variants are often also included on these platforms.

The large, population-based genetic dataset available in the Utah Suicide Genetic Risk Study (USGRS) provides unique opportunities to uncover both common and rare variants leading to risk of suicide death. Genetic data from 4,382 suicide death cases were generated with the Illumina PsychArray BeadChip, a high-density, genome-wide microarray developed specifically for large-scale genetic studies of psychiatric disorders. The PsychArray includes 265,000 common tag-SNPs as well as 245,000 markers from Illumina’s Exome BeadChip, developed to capture coding variants that alter gene function as well as 50,000 additional markers associated with psychiatric conditions (https://www.illumina.com/products/by-type/microarray-kits/infinium-psycharray.html). These latter markers were selected specifically for their potential functional impact and relevance to psychiatric disease.

Modern GWAS focus on the detection and analysis of common genetic risk factors, and SNPs with MAF <1%, such as the rare variants included on the PsychArray, are traditionally removed as being less informative for this study design. However, the removal of content specifically chosen for both functional and disease relevance represents a missed opportunity for the discovery of rare, more highly penetrant risk factors for complex psychiatric traits. There has been no large-scale effort to evaluate the PsychArray’s rare variant content in the context of complex psychiatric traits. Due to the popularity of the PsychArray as a genotyping platform, content traditionally not considered in GWAS is an untapped resource to investigate rare variants involved in suicide, as well as other complex psychiatric traits. This analysis requires special methodological considerations, including rigorous attention to quality control issues followed by comparison with external control data as well as secondary molecular validation of variants of interest.

Here, we investigate the role of rare variants in suicide death. First, we identified rare variant content on the PsychArray that is significantly associated with suicide death using a case-control design, comparing USGRS cases against controls from the large gnomAD resource. Next, we prioritized rare variants by comparing with an independent control cohort genotyped on the PsyschArray, and validated these variants in the USGRS with sequencing data. Follow-up analyses of these variants identified important mechanistic pathways and cross-trait associations. Finally, we characterized the phenotypic attributes of all suicide cases with these rare variants.

## MATERIALS AND METHODS

### Utah Suicide Genetic Research Study Cohort

In Utah, suicide rates have risen steadily for the past 20 years, and Utah currently has the sixth highest suicide rate in the nation.^17^ The USGRS has a biospecimen resource of 6,202 population-ascertained DNA samples from suicide decedents (4,837 males and 1,365 females). This resource grows by ∼650 cases per year reflective of the rate of suicide in Utah. Samples are ascertained through a two-decade collaboration with the centralized Utah Office of the Medical Examiner (OME), ensuring consistency of suicide determination, and facilitating ongoing state-wide collection of 500-600 cases per year. Since 1998, we have collected de-identified DNA samples from suicide deaths with Institutional Review Board (IRB) approval. DNA is extracted from blood using the Qiagen Autopure LS automated DNA extractor (www.qiagen.com). OME cases are then linked to the Utah Population Database (UPDB), which includes multi-generational genealogical, demographic, geographic and medical information for over 11 million individuals, using secure computer servers. After linkage, suicide cases are stripped of identifying information before transfer to the research team. All subjects are referenced by anonymous IDs and no contact is possible with living family members. The state-wide sample ascertainment is population-based, and not limited to cases within a research or clinical cohort or with any psychiatric diagnosis.

### PsychArray Genotyping and Quality Control

Genetic data from 4,379 suicide cases were generated with the Illumina PsychArray BeadChip. This is a high-density, genome-wide microarray developed specifically for large-scale genetic studies of psychiatric conditions. Genotyping was performed in 6 batches with duplicate samples across batches to allow for quantification and modeling of any batch effects. Quality control of all PsychArray genotypes was performed in GenomeStudio for each batch separately. SNPs were retained if the GenTrain score was > 0.5 and the Cluster separation score was > 0.4. SNPs were converted to HG19 plus strand format then batches were combined. PLINK^18,19^ was used to remove duplicated SNPs or individuals, poorly genotyped cases, monomorphic SNPs, and non-autosomal SNPs. A total of 368,152 SNPs were retained after quality control.

### Ancestry Estimation

Ancestry estimates were computed using ADMIXTURE.^20^ PsychArray genotypes were compared to 1000 Genomes (https://www.internationalgenome.org/data/) reference panel. Ten percent of Utah suicide cases are not of predominate European ancestry (e.g., Latinx, Asian, African, or of mixed ancestries). Rare variants are expected to display stronger patterns of population stratification than common variants.^21^ To control for possible allele frequency differences due to ancestry, we confined our analyses to the 2,735 cases that had ancestry estimates of at least 90% Non-Finnish European (NFE) ancestry (Supplementary Figure 1). This is a conservative estimate of ancestry as most samples in the USGRS are predominately European. 85% of the USGRS suicide cases are >75% NFE ancestry.

### Sample Relatedness

Estimates of pairwise identity by descent (IBD) were calculated using PLINK. Pairs of related individuals (third degree or closer) were identified with pi-hat values greater than 0.12. One member of each of the 63 identified pairs of relatives was randomly removed from the NFE subset and 2,672 remaining individuals were assessed.

### Control Data

Control data were downloaded from the Genome Aggregation Database (gnomAD v2.1).^22^ gnomAD provides allele frequencies from aggregated genomic data from sequencing studies of over 141,000 unrelated individuals. The NFE gnomAD subset excluding cases with neurological phenotypes, referred to as non-neuro (NN), was used in this study. This NFE-NN subset provides a comparison group eliminating possibly confounding cases with neuropsychiatric conditions. Genome and exome VCFs from the gnomAD data were filtered using bcftools^23^ to keep only SNPs at genomic sites that passed all gnomAD quality filters (see https://macarthurlab.org/2018/10/17/gnomad-v2-1/). Genomic sites identified as multiallelic in gnomAD were also removed. Biallelic SNPs present in both genome and exome datasets and passing QC filters were combined. SNPs with MAF <0.01 (2,813,863 sites) in the combined NFE-NN dataset were matched to filtered PsychArray SNPs, resulting in a set of 36,215 SNPs in common between the two datasets.

### Variant Annotation

Before proceeding with association tests, the SNPs in both the PsychArray and gnomAD data were prioritized for functional relevance by annotating all SNPs using Combined Annotation Dependent Depletion (CADD).^24^ CADD prioritizes functional, deleterious and pathogenic coding and non-coding variants across many functional annotations. If variants had at least one of the following CADD annotations they were retained in the analysis: 1) probably or possibly damaging in PolyPhen; 2) deleterious in SIFT or 3) CADD scaled C-score (phred) >=10 (a score reflecting a deleterious ranking of the variant compared to all possible substitutions in the genome). Higher CADD phred scores indicate an increased likelihood that a variant will have meaningful functional consequences. For example higher CADD phred scores are associated with larger negative effects of rare variants on serum urate.^25^ In total, 30,377 SNPs were retained after CADD filtering.

### Single Variant Analyses

Allele counts in suicide cases and gnomAD controls were compared using Fisher’s exact test. Genomic sites having significant p-values after Bonferroni correction for 30,377 multiple comparisons were retained for stage 2 analyses (see Figure 1). We chose this conservative correction method assuming independence of all comparisons. After this first stage, 27 variants survived multiple testing. Next, in a stage 2 analysis, we performed an extension of the gnomAD comparisons using an independent control dataset genotyped on the PsychArray. Stage 2 considered only significant results from the gnomAD comparisons, comparing the frequencies of 27 variants in the USGRS suicides to an independent cohort of 21,324 NFE controls without psychiatric conditions generated by the Psychiatric Genomics Consortium (PGC). SNP allele counts in suicide cases and PGC control individuals were compared using a Fisher’s exact test using a replication significance threshold of P <2.17E-03. Genomic sites having significant p-values after Bonferroni correction were retained for stage 3 validation analysis.

**Figure 1:**
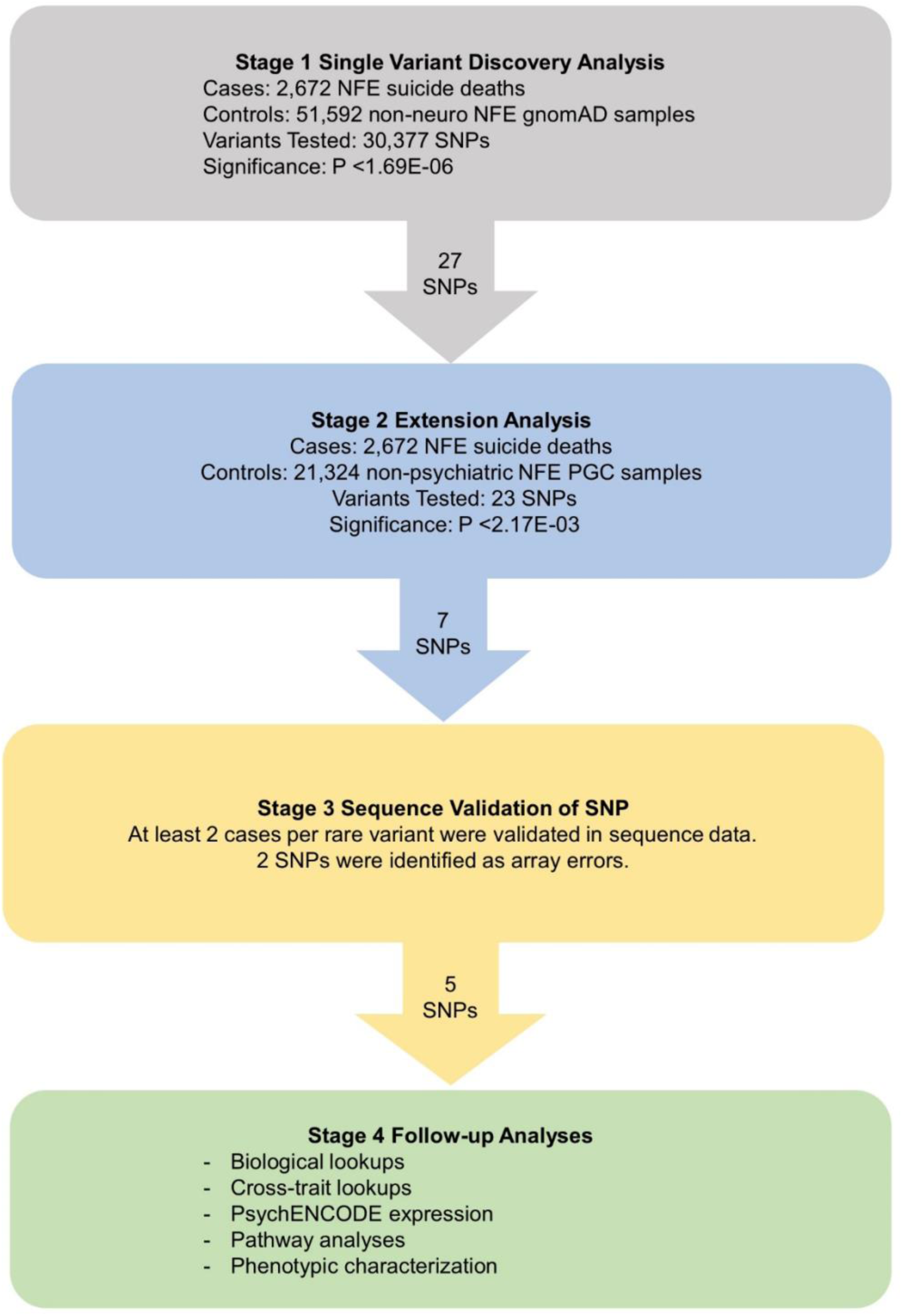
Study design and workflow diagram for rare variant analyses. gnomAD, Genome Aggregation Database v2.1; non-neuro, individuals who were not ascertained for having a neurological condition in a neurological case/control study; NFE, non-Finish European.

### Validation of Rare Variant Content

A subset of 286 suicide cases with PsychArray data also have whole genome sequence (WGS) data generated with Illumina technology. USGRS WGS data was jointly processed using a BWA/GATK-Haplotype caller sequence analysis pipeline^26^ with control WGS data, including unrelated cases from the 1000 Genomes CEU cohort, 1000 Genomes subjects from Great Britain (matching in ancestry to Utah cases), and a control group of Utah elderly (>age 90) healthy subjects. USGRS WGS data had mean coverage of 30x-60x. Data were assessed for duplicate rate, fraction of properly paired reads, absolute and relative number of calls for different variant classes, and the Ts/Tv ratio within coding sequences. This sequence data allowed for direct validation of rare variants where cases had both array and sequence data. If cases carrying alternate alleles surviving stages 1 and 2 were not present in this WGS dataset but were significant in stage 1 and 2 analyses, we directly validated the variant using Sanger sequencing. Only variants where presence of the allele was validated in sequence data were considered.

### Phenotyping

All cases in the USGRS have demographic information and nearly all cases have cause of death from Vital Records data and Medical Examiner autopsy reports. Additionally, approximately 80% of cases have electronic medical record (EMR) data, specifically International Classification of Diseases (ICD) diagnostic codes (ICD-9/ICD-10). Missing EMR data can occur for many reasons (e.g., individuals not seeking medical attention, care received out of the period of UPDB record coverage). The phenotypic characteristics of the 2,672 individuals used in the rare variant analyses were described by age, sex, cause of death and 30 relevant co-occurring psychiatric and related medical phenotypes established by hierarchically aggregating ICD codes in EMR data using the ICD system and with review by clinical experts on our team (A.D., B.K., and E.M.) (Supplementary Table 1). Counts of occurrences of EMR phenotypes of cases with and without identified risk SNPs were compared using Fisher’s exact test with a Bonferroni significance threshold of P <0.0017.

### Follow-Up Analyses

Clinically relevant trait associations were assessed at both the SNP and gene level. Identified SNPs that were significant in all three stages were checked against the ClinVar^27^, dbSNP^28^ and DisGeNET^29^ databases. At the gene level, trait associations were assessed by a search of the literature, using the NHGRI-EBI GWAS Catalog^30^, and PsychENCODE differential gene expression.^31^ Pathway analyses were also conducted at the gene level using FUMA GENE2FUNC software^32^ to determine pathway enrichments in the Gene Ontology (GO) gene sets compared to all genes in the genome. We conducted tests within genes meeting significance and validation in this study, and additional tests including genes implicated by GWAS studies on suicidal behavior. 102 genes from GWAS studies focusing on discovery of common variants associated with suicide attempt and death that were prioritized by *Gonzalez-Castro et al*.^33^ were used in the pathway analyses. P-values for enrichment analyses were adjusted for multiple testing using the Benjamini-Hochberg false discovery rate (FDR) method.

## RESULTS

### Single Variant Analyses

We conducted a single variant association analysis of 30,377 rare putatively functional SNPs present on the PsychArray genotyping array in 2,672 USGRS suicides of NFE ancestry and 51,583 publicly available NFE controls from gnomAD. The average CADD PHRED score for the 30,377 variants tested was 21.54 (5.96 SD). 2,528 SNPs had nominally significantly elevated allele frequencies in suicide cases compared with gnomAD NFE-NN controls. The average CADD PHRED score for the 2,528 nominally significant variants was 22.05 (6.25 SD). In our primary (stage 1) analysis, we identified 27 SNPs on the PsychArray with predicted functional consequences with significantly elevated allele frequencies after Bonferroni correction in suicide cases compared with gnomAD NFE-NN controls (Supplementary Table 2). We chose this conservative correction method due to possible array heterogeneity and estimated increased error rates in genotyping rare variants.^12^

To control for differences in genotyping platform we performed stage 2 extension analyses with an independent PsychArray control set. Stage 2 considered only significant results from the stage 1 comparisons. 23 of the 27 significant SNPs from stage 1 were comparable in the stage 2 PGC control set comprised of 21,324 NFE controls without psychiatric conditions. Four of the 27 SNPs were not comparable as they did not pass QC thresholds. Seven of the 23 tested variants had significantly elevated allele frequencies in suicide cases in stage 2 when compared with controls after Bonferroni correction (Supplementary Table 3).

In stage 3 analyses, we validated the genotype call from the PsychArray genotyping platform. Two of the seven variants did not validate in sequence data. Five novel, high-impact, rare SNPs (MAF <1%) survived all stages of our study (Table 1; Figure 2). All five variants had CADD PHRED scores greater than 22 (Table 2) indicative of the top 1% of deleterious substitutions in the human genome.^24^ These risk variants include protein coding variants with high predicted deleterious impact in *SNAPC1* (rs75418419), *TNKS1BP1* (rs143883793), *ADGRF5* (rs149197213), *PER1* (rs145053802) and *ESS2* (rs62223875). Together, these five rare variants were present in 119 of the 2,672 suicide cases of NFE ancestry (4.45% of cases). Figure 2 shows odds ratios and confidence intervals for each variant.

**Table 1.** Rare missense variants associated with suicide death. *a* minor allele frequency in gnomAD v2.1 non-Finnish-European, non-neuro subset; *b* minor allele frequency in non-Finnish-European suicide cases; *c* p-value <1.7E-6 (Bonferroni correction for 30,377 tests); *d* one individual was homozygous for the alternate allele.

| SNP location<br>(GRCh37) | Reference<br>allele | Alternate<br>allele | PsychArray<br>SNP | rs# | Gene Name | CADD<br>PHRED<br>score | gnomAD<br>MAF<sub>a</sub> | Suicide<br>MAF<sub>b</sub> | gnomAD<br>reference<br>allele count | gnomAD<br>alternate<br>allele count | Suicide<br>reference<br>allele<br>count | Suicide<br>alternate<br>allele<br>count | P-value<sub>c</sub> | # suicide<br>cases |
| --- | --- | --- | --- | --- | --- | --- | --- | --- | --- | --- | --- | --- | --- | --- |
| 17:8045136 | G | A | exm1290985 | rs145053802 | *PER1* | 23.2 | 4.48E-04 | 3.56E-03 | 102634 | 46 | 5325 | 19 | 2.06E-10 | 19 |
| 6:46851334 | C | T | exm553084 | rs149197213 | *ADGRF5 (GPR116)* | 24.7 | 8.82E-04 | 4.68E-03 | 101994 | 90 | 5317 | 25 | 3.94E-10 | 25 |
| 11:57070069 | C | T | exm910447 | rs143883793 | *TNKS1BP1* | 23.8 | 1.58E-03 | 5.43E-03 | 102618 | 162 | 5313 | 29 | 8.14E-08 | 28<sub>d</sub> |
| 14:62244854 | C | T | exm1106246 | rs75418419 | *SNAPC1* | 22.3 | 2.05E-03 | 6.18E-03 | 101432 | 208 | 5311 | 33 | 1.74E-07 | 33 |
| 22:19127403 | C | T | exm1584742 | rs62223875 | *ESS2 (DGCR14)* | 24.7 | 4.75E-04 | 2.81E-03 | 103117 | 49 | 5327 | 15 | 3.80E-07 | 15 |

**Table 2.** Descriptive data for suicide cases with and without five identified risk SNPs.

|  | cases without risk SNPs | cases with risk SNPs |
| --- | --- | --- |
| # Individuals | 2553 | 119 |
| Mean age (SD) in years | 41.42 (17.63) | 40.71 (17.74) |
| Male | 0.79 | 0.82 |
| Female | 0.21 | 0.18 |
| With method of death data | 0.96 | 0.94 |
| Firearm | 0.54 | 0.53 |
| Asphyxiation | 0.30 | 0.30 |
| Overdose/poison | 0.12 | 0.13 |
| Violent trauma | 0.03 | 0.03 |
| Other method of death | 0.01 | 0.01 |

**Figure 2:**
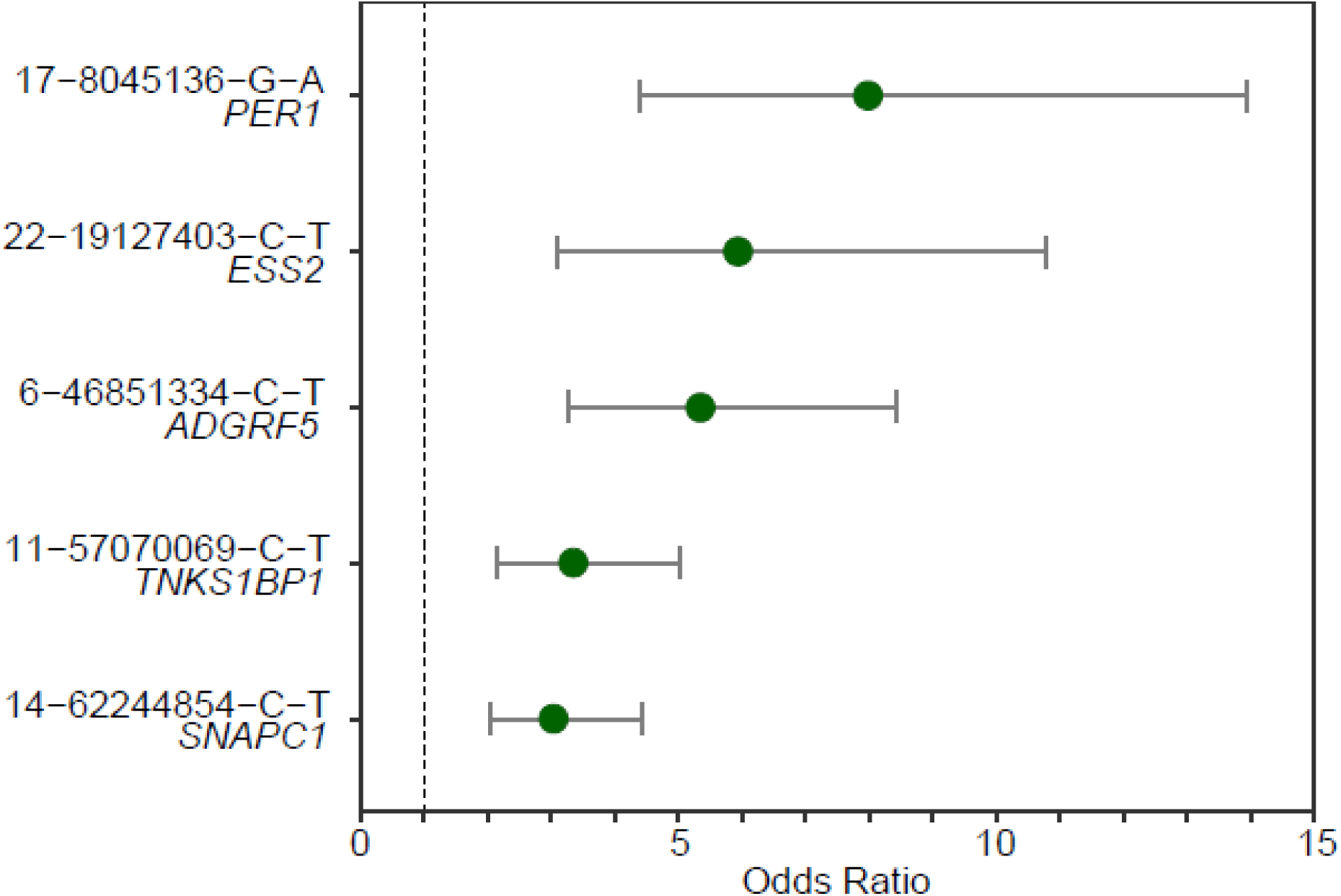
Estimated odds ratios and 95% confidence intervals of significant variants in suicide cases compared to PGC controls. Dashed line is placed at 1. Variant labels are in the format chromosome, position, reference allele, alternate allele.

### Trait Associations, Gene Function, Expression and Pathway Analysis

At the gene level, two of the identified genes with rare risk variants had prior associations with suicidal behavior (*PER1*^34,35^ and *SNAPC1*^36^), and three had prior associations with severe psychopathology (*PER1, TNKS1BP1*, and *ESS2*) (see discussion for details). No prior trait associations were found in the literature or in searched databases for the five specific SNPs identified in this study.

*PER1, TNK1S1BP1*, and *ADGRF5* showed significant enrichments in GO biological process gene sets with the 102 prioritized genes from genome-wide studies of suicidal behavior (Supplementary Table 4). *TNKS1BP1* and *ADGRF5* were significant in the Homeostatic Process gene set (M11223), together with *PDE4B, DISC1, ITGB1, TMX3, PRKCE, SLC4A4*, and *IL7. PER1* is a member of the Regulation of Intracellular Signal Transduction (M11395) pathway - together with *ITGB1, FGD4, SIPA1L1, CDH13, PRKCE, LRRTM4, SH3RF1, KIAA1244, NTRK2. ADGRF5 also* implicated the Regulation of Immune System Process (M13496) pathway, together with *PDE4B, CAMK1D, ITGB1, CYP19A1, CD300LB, PRKCE, RBFOX2*, and *IL7*. FUMA pathway enrichment analyses using only the five final genes remaining after stage 3 analysis did not implicate any biological pathways.

### Phenotypic Associations

Demographic and diagnostic data were compared between the NFE suicide cases without rare risk variants and the suicide cases with identified rare risk variants (Table 2, Table 3). Use of firearms was the most common method of death for suicide cases with and without rare variants. There was not a significant difference in the age of cases with and without risk SNPs (t= 0.4293, p = 0.6677). None of the 30 EMR diagnostic categories differed significantly (Bonferroni significant threshold p < 0.00167) between suicide cases with and without identified risk SNPs (Table 3).

**Table 3.** Prevalence of Broad EMR diagnostic categories for suicide cases with and without five identified risk SNPs.

| EMR Phenotype | Cases Without Risk SNPs | Cases with Risk SNPs |
| --- | --- | --- |
| Accidental Trauma | 0.634 | 0.604 |
| Alcohol | 0.246 | 0.250 |
| Anxiety Non Trauma | 0.366 | 0.344 |
| Asthma | 0.112 | 0.125 |
| Bipolar | 0.128 | 0.083 |
| Cardiovascular | 0.458 | 0.406 |
| COPD | 0.073 | 0.052 |
| Dementia Neurodegenerative | 0.117 | 0.125 |
| Depression | 0.497 | 0.469 |
| Developmental | 0.016 | 0.031 |
| Drug | 0.275 | 0.292 |
| Eating | 0.011 | 0 |
| Head Injury | 0.066 | 0.083 |
| Immune Autoimmune | 0.193 | 0.271 |
| Impulse Control Disorders | 0.093 | 0.042 |
| Injury By Law Enforcement | 0.004 | 0 |
| Interpersonal Trauma | 0.345 | 0.365 |
| Migraine | 0.090 | 0.125 |
| Nutrition Metabolism | 0.415 | 0.365 |
| Obesity | 0.208 | 0.250 |
| Pain | 0.681 | 0.656 |
| PD Cluster A | 0.003 | 0 |
| PD Cluster B | 0.093 | 0.063 |
| PD Cluster C | 0.011 | 0.000 |
| Schizophrenia Schizoaffective | 0.038 | 0.052 |
| Seizure | 0.040 | 0.031 |
| Sleep | 0.258 | 0.302 |
| Somatic | 0.093 | 0.104 |
| Suicidal Ideation | 0.163 | 0.125 |
| Self Injury | 0.212 | 0.198 |

For cases carrying the rare risk variant at the five SNPs who also had linked EMR data, at least 50% of individuals had chronic/acute pain diagnoses (Supplementary Table 5). For cases with identified risk variants in *TNKS1BP1, ESS2, ADGRF5* or *PER1* and EMR data, >45% of cases had depression diagnoses. For cases with identified risk variants in *SNAPC1, TNKS1BP1, ESS2* or *PER1* and EMR data, >55% of cases had diagnoses related to accidental trauma. However, these frequencies are similar when compared against suicide cases without the risk variants.

Of the 15 suicide cases with the rare missense variant in *ESS2*, 14 had EMR data. 12 of 14 of these cases had ICD diagnosis codes indicative of accidental trauma. One case had rare missense variants in both *SNAPC1* and *ADGRF5.* This case was 16 years old at the time of death. Only 1 ICD code was observed for this individual (ICD9 787.03 “vomiting alone”).

Two cases with the identified risk variant in *SNAPC1* had bipolar diagnoses. For risk SNP carriers in *TNKS1BP1*, one case had schizophrenia diagnoses, one case had both bipolar and schizophrenia diagnoses and one case had bipolar and schizoaffective diagnoses. One case had schizoaffective and bipolar disorder diagnoses and the risk SNP in *PER1*. For *ESS2* one case had schizoaffective disorder diagnoses and for *ADGRF5* one case had bipolar disorder diagnoses in EMR data.

## DISCUSSION

This study identified five novel variants likely to impact gene function that are significantly associated with suicide death. These variants implicate the following genes: *SNAPC1, TNKS1BP1, ADGRF5, PER1*, and *ESS2.* Our study included rigorous thresholds for quality control and ancestry, and all surviving variants were validated in sequence data, addressing concerns of potential error in PsychArray rare variant content. Of note, the significant variants in our study are closer to 1% in frequency; variants that were <0.1% did not validate in sequence data.

At the gene level, our analyses revealed two genes with prior evidence linking them to suicide risk: *PER1* and *SNAPC1*, though specific functional risk variants were not identified in those previous studies. *PER1* is a circadian clock gene that plays an essential role in generating circadian rhythms.^37^ *PER1* has been implicated in long-term memory formation^38^, response to insufficient sleep^39^ and regulation of neuroinflammation.^40^ Altered expression of *PER1* has been found in individuals with schizophrenia^41,42^ and *PER1* knock-out mice exhibit ADHD-like symptoms and reduced levels of dopamine.^43^ *PER1* was significantly downregulated in the prefrontal cortex of individuals who died by suicide^34^ and was differentially expressed in suicide attempters compared with controls.^44^ Postmortem brain tissue expression results from the psychENCODE sample showed *PER1* has increased expression in schizophrenia (FDR p=0.0006) and bipolar cases (FDR p=0.0016).

*SNAPC1* encodes a subunit of the Small Nuclear RNA Activating Complex (SNAPc), which activates RNA polymerase II and III and is required for translation of small nuclear RNA.^45,46^ *SNAPC1* was also identified in a genome-wide significant region in extended Utah families enriched for suicide deaths.^47^

The other genes impacted by rare variants in our study have been implicated in psychiatric disorders or immune related phenotypes. *TNKS1BP1* plays a role in DNA damage repair, genome stability, and regulation of the actin cytoskeleton.^48–50^ Expression results from psychENCODE showed increased expression differences for *TNKS1BP1* in schizophrenia (FDR p=0.02). *ADGRF5*, formally *GPR116*, encodes a G protein-coupled receptor that plays a critical role in regulating pulmonary immune response.^51,52^ The GWAS catalog identified *ADGRF5* as having associations with four blood related traits (blood protein levels, white blood cell count, mean platelet volume, monocyte count) and adolescent idiopathic scoliosis. *ESS2* (formally known as *DGCR14*) is a nuclear protein that enhances the transcriptional activity of the retinoic acid receptor, and is a component of the spliceosome C complex which removes introns from pre-mRNA^53^. Mutations in *ESS2* are associated with risk for schizophrenia. ^54,55^ In addition, deletions in *ESS2* are associated with DiGeorge syndrome, which causes immunodeficiency, heart defects, skeletal abnormalities, psychiatric disturbance, and alterations in REM sleep.^55^

The associations of *AGDRF5* with pulmonary immune response, blood phenotypes, and scoliosis and the association of *PER1* with neuroinflammation may reflect recent data suggesting the importance of immune response and inflammation in risk generally of psychopathology^56^ and specifically of suicide.^57^ Supporting this association, *ADGRF5*, with genes implicated by GWAS studies of suicidal behavior showed an enrichment in immune pathways.^33^

The 119 suicide cases with these variants represent 4.45% of the European genotyped study sample that was the focus of this study. 96 of the 119 cases (81%) had EMR data and these cases showed evidence of chronic pain, depression, and trauma phenotypes in electronic medical records data. No clear differences with regards to medical diagnoses from EMR were noted between suicide cases with and without the 5 identified risk SNPs, although as expected for rare variants, we were underpowered to detect differences if they did exist. While previously published associations suggested risk of schizophrenia for *PER1, ESS2* and *TNKS1BP1*, only 10% of cases with risk alleles in these genes had schizophrenia or schizoaffective diagnoses, vs 4% of cases without these rare variants. *PER1* has also previously been associated with bipolar disorder, yet only 1 of the 15 cases with the risk SNP and EMR data had a bipolar diagnosis.

Our analysis presents an efficient, cost-effective study of putative functional variants in the Utah Suicide Genetic Research Study (USGRS), a research resource that includes a large sample of molecular data from suicide deaths linked to extensive demographic and medical record data. Importantly, most large studies of the genetics of suicide focus on suicidal behaviors, which are relatively common (4.3% per year^58^). However, suicidal behaviors imperfectly predict the more extreme outcome of suicide death (0.01-0.02% per year^58^). The USGRS provides a resource for genetic discovery of risks specifically associated with the outcome of suicide death. Every discovery of specific genetic risk factor associated with suicide death will bring us closer to the goal of targeted prevention for those at risk. This study complements our recent genome-wide association study^59^, which focuses on the discovery of common variation associated with suicide death.

This study demonstrates an important role for rare variants in the polygenic architecture of suicide risk. Importantly, rare variants are more functionally tractable than common variants and may reveal unique insights into the mechanisms of risk. We have made careful use of rare, functional content on the genome-wide array to reach the discoveries reported in this study.

## LIMITATIONS

This study has two important limitations. First, while taking a conservative approach and refining our data to pre-selected, rare array content with previous evidence of relevance to psychiatry, these data represent only a tiny proportion of the potential rare, functional risk variation in the genome. Second, because rare variants are particularly susceptible to ancestry effects^16^, we confined our analysis to individuals of European ancestry, the group making up the largest proportion of our data resource. This may limit generalizability of findings to other ancestry groups.

In spite of these limitations, our results represent important novel risk variants and genes associated with risk for suicide death. Our success with this approach may in part reflect the design of the PsychArray, providing highly relevant content. Success may also be driven by our analysis of suicide death, an extreme phenotypic outcome.

## CONCLUSION

Overall, the research community has made great progress with genetic discovery of both common and rare variants leading to risk of complex phenotypes. The aggregated effects of common and rare variants are starting to make a difference for individuals with these conditions. While the genetic study of suicide death has far to go, as discoveries accumulate, aggregation of results into larger risk pathways will begin to have an impact on personalized risk prediction and prevention.

## Supporting information

Supplementary Material PGC author list

## ACKNOWLEDGEMENTS

This work was supported by: the American Foundation for Suicide Prevention (ED, AVB), the National Institute of Mental Health, R01MH099134 (HC) and K01MH093731 (AD), the Brain & Behavior Research Foundation Young Investigator Award (AD), the Clark Tanner Foundation (HC, TD, AVB). Processing of samples was done with assistance from GCRC M01-RR025764 from the National Center for Research Resources. Partial support for all data sets within the UPDB was provided by the University of Utah Huntsman Cancer Institute. We especially thank the University of Utah, the Psychiatric Genomics Consortium and OME staff whose hours of work made this study possible.

## SUPPLEMENTARY MATERIAL LEGENDS

### SUPPLEMENTARY TABLES

**Table S1.** ICD code groupings for EMR data.

**Table S2.** Details of 27 PsychArray SNPs that were identified in stage 1 of study. *a* minor allele frequency in gnomAD v2.1 non-Finnish-European, non-neuro subset; *b* minor allele frequency in NFE suicide cases; *c* major allele count for gnomAD controls; *d* minor allele count for gnomAD controls; *e* major allele count for suicide cases; *f* minor allele count for suicide cases; *g* Fisher’s exact test p-value, less than Bonferroni correction for 30,377 tests; *h* If variant was present in WGS dataset of 286 suicide cases.

**Table S3.** Results of stage 2 extension analyses. *a* minor allele frequency in Psychiatric Genomics Consortium NFE controls without psychiatric conditions *b* minor allele frequency in NFE suicide cases; *c* major allele count for PGC controls; *d* minor allele count for PGC controls; *e* major allele count for suicide cases; *f* minor allele count for suicide cases; *g* Fisher’s exact test p-value, less than Bonferroni correction for 23 tests; shaded rows indicate SNPs that were significant in stage 1 but did not survive QC thresholds in stage 2 comparisons.

**Table S4.** Gene set enrichment gene list and results.

**Table S5.** Prevalence of 30 EMR phenotypes in cases carrying minor allele of rare risk variants.

### SUPPLEMENTARY FIGURES

**Figure S1.** (Left) The first two PCs plotted for suicide genotypes (gray) and 1KGP super population reference data. Rare variants tend to be population specific. Therefore, we included only non-Finnish European (NFE) individuals in our analyses, indicated within the red square. (Right). The first two PCs plotted for samples used in the analyses, including NFE suicide samples (gray) and 1KGP population reference data.

