## Supplementary Table 2 for "Rare protein coding variants implicate genes involved in risk of suicide death"

**Table S2.** Details of 27 PsychArray SNPs that were identified in stage 1 of study. *a* minor allele frequency in gnomAD v2.1 non-Finnish-European, non-neuro subset; *b* minor allele frequency in NFE suicide cases; *c* major allele count for gnomAD controls; *d* minor allele count for gnomAD controls; *e* major allele count for suicide cases; *f* minor allele count for suicide cases; *g* Fisher’s exact test p-value, less than Bonferroni correction for 30,377 tests; *h* If variant was present in WGS dataset of 286 suicide cases.

| SNP Name | PsychArray SNP | GeneName | gnomAD MAF^a^ | SUI  MAF^b^ | gnomAD_c2^c^ | gnomAD_C1^d^ | SUI_C2^e^ | SUI_C1^f^ | Pvalue^g^ | In seq data^h^ | # suicide cases |
| --- | --- | --- | --- | --- | --- | --- | --- | --- | --- | --- | --- |
| 1:12837720:G:A | exm17166 | PRAMEF12 | 5.81E-03 | 3.03E-02 | 98351 | 575 | 5155 | 161 | 5.47E-56 | Y | 161 |
| 21:15599354:G:A | exm1562529 | RBM11 | 7.24E-03 | 1.80E-02 | 99506 | 726 | 5248 | 96 | 4.84E-14 | Y | 95 (1 homozygote) |
| 3:50609625:C:T | exm318388 | HEMK1 | 1.95E-05 | 1.52E-02 | 102408 | 2 | 5253 | 81 | 3.19E-103 | N | 81 |
| 19:49671207:G:A | exm1489934 | TRPM4 | 1.64E-03 | 1.39E-02 | 99009 | 163 | 5270 | 74 | 1.87E-37 | Y | 74 |
| 2:46707827:T:C | exm191574 | TMEM247 | 7.27E-04 | 1.31E-02 | 59119 | 43 | 5262 | 70 | 9.03E-47 | Y | 70 |
| 11:96117537:A:C | exm949941 | CCDC82 | 3.08E-03 | 8.63E-03 | 101748 | 314 | 5284 | 46 | 5.72E-09 | Y | 46 |
| 7:87032531:T:C | exm631571 | ABCB4 | 3.88E-05 | 7.93E-03 | 103168 | 4 | 5252 | 42 | 9.50E-51 | N | 42 |
| 14:62244854:C:T | exm1106246 | SNAPC1 | 2.05E-03 | 6.18E-03 | 101432 | 208 | 5311 | 33 | 1.74E-07 | Y | 33 |
| 13:48528645:G:A | newrs121908538 | SUCLA2 | 3.88E-05 | 5.99E-03 | 103054 | 4 | 5312 | 32 | 6.57E-38 | N | 32 |
| 15:42379900:T:C | exm1153294 | PLA2G4D | 2.43E-04 | 5.61E-03 | 102821 | 25 | 5314 | 30 | 5.46E-25 | Y* (tags indel) | 30 |
| 11:57070069:C:T | exm910447 | TNKS1BP1 | 1.58E-03 | 5.43E-03 | 102618 | 162 | 5313 | 29 | 8.14E-08 | Y | 28 (1 homozygote) |
| 11:70170536:A:G | exm936578 | PPFIA1 | 6.80E-05 | 5.45E-03 | 102945 | 7 | 5295 | 29 | 6.34E-32 | N | 29 |
| 6:46851334:C:T | exm553084 | ADGRF5 (GPR116) | 8.82E-04 | 4.68E-03 | 101994 | 90 | 5317 | 25 | 3.94E-10 | Y | 25 |
| 17:8045136:G:A | exm1290985 | PER1 | 4.48E-04 | 3.56E-03 | 102634 | 46 | 5325 | 19 | 2.06E-10 | Y | 19 |
| 4:76956246:T:C | exm407053 | CXCL11 | 2.91E-04 | 3.00E-03 | 103002 | 30 | 5326 | 16 | 2.86E-10 | Y* (tags indel) | 16 |
| 22:19127403:C:T | exm1584742 | ESS2 | 4.75E-04 | 2.81E-03 | 103117 | 49 | 5327 | 15 | 3.80E-07 | Y | 15 |
| 19:55143609:G:A | variant.58198 | LILRB1 | 0.00E+00 | 2.07E-03 | 103160 | 0 | 5297 | 11 | 3.82E-15 | N | 11 |
| 19:6427409:C:T | exm1413868 | SLC25A41 | 1.99E-05 | 1.88E-03 | 100516 | 2 | 5316 | 10 | 6.21E-12 | N | 10 |
| 3:9988425:T:C | exm289168 | PRRT3 | 8.92E-05 | 1.88E-03 | 67280 | 6 | 5306 | 10 | 2.33E-08 | N | 10 |
| 20:52789469:T:C | exm1551406 | CYP24A1 | 7.76E-05 | 1.87E-03 | 103128 | 8 | 5334 | 10 | 2.54E-09 | N | 10 |
| 19:49351155:A:G | exm1488444 | PLEKHA4 | 0.00E+00 | 1.71E-03 | 85282 | 0 | 5267 | 9 | 7.68E-12 | N | 9 |
| 18:44221971:C:T | exm1385333 | LOXHD1 | 3.12E-05 | 1.31E-03 | 64020 | 2 | 5321 | 7 | 4.93E-07 | N | 7 |
| 1:230979494:G:A | exm158330 | C1orf198 | 1.95E-05 | 1.31E-03 | 102738 | 2 | 5327 | 7 | 2.34E-08 | N | 7 |
| 13:25479542:C:G | exm1058471 | CENPJ | 0.00E+00 | 1.15E-03 | 103158 | 0 | 5232 | 6 | 1.27E-08 | N | 6 |
| 15:38643456:T:C | exm1147382 | SPRED1 | 1.94E-05 | 1.13E-03 | 103148 | 2 | 5308 | 6 | 3.54E-07 | N | 6 |
| 17:2227988:T:C | exm2225932 | TSR1 | 0.00E+00 | 9.41E-04 | 102122 | 0 | 5309 | 5 | 2.96E-07 | N | 5 |
| 7:63982623:T:C | exm623076 | ZNF680 | 0.00E+00 | 9.41E-04 | 102156 | 0 | 5307 | 5 | 2.95E-07 | N | 5 |
