## Supplementary Table 3 for "Rare protein coding variants implicate genes involved in risk of suicide death"

**Table S3.** Results of stage 2 extension analyses. *a* minor allele frequency in Psychiatric Genomics Consortium NFE controls without psychiatric conditions *b* minor allele frequency in NFE suicide cases; *c* major allele count for PGC controls; *d* minor allele count for PGC controls; *e* major allele count for suicide cases; *f* minor allele count for suicide cases; *g* Fisher’s exact test p-value, less than Bonferroni correction for 23 tests; shaded rows indicate SNPs that were significant in stage 1 but did not survive QC thresholds in stage 2 comparisons.

| SNP Name | PsychArray SNP | GeneName | PGC  MAF^a^ | SUI  MAF^b^ | PGC_C2^c^ | PGC_C1^d^ | SUI_C2^e^ | SUI_C1^f^ | Pvalue^g^ |
| --- | --- | --- | --- | --- | --- | --- | --- | --- | --- |
| 1:12837720:G:A | exm17166 | PRAMEF12 | 0.03574 | 0.03029 | 38432 | 1474 | 5155 | 161 | 0.01373 |
| 21:15599354:G:A | exm1562529 | RBM11 | 0.01651 | 0.01796 | 40046 | 677 | 5248 | 96 | 0.4617 |
| 3:50609625:C:T | exm318388 | HEMK1 | 0.01408 | 0.01519 | 40224 | 578 | 5253 | 81 | 0.5397 |
| 19:49671207:G:A | exm1489934 | TRPM4 | 0.01732 | 0.01385 | 39924 | 715 | 5270 | 74 | 0.04966 |
| 2:46707827:T:C | exm191574 | TMEM247 | 0.01238 | 0.01313 | 40376 | 512 | 5262 | 70 | 0.6952 |
| 11:96117537:A:C | exm949941 | CCDC82 | 0.008546 | 0.00863 | 40618 | 353 | 5284 | 46 | 1 |
| 7:87032531:T:C | exm631571 | ABCB4 | 0.01415 | 0.007934 | 40202 | 588 | 5252 | 42 | 5.36E-05 |
| 14:62244854:C:T | exm1106246 | SNAPC1 | 0.002271 | 0.006175 | 41202 | 93 | 5311 | 33 | 4.00E-06 |
| 13:48528645:G:A | newrs121908538 | SUCLA2 | NA | 0.005988 | NA | NA | 5312 | 32 | NA |
| 15:42379900:T:C | exm1153294 | PLA2G4D | 0.005665 | 0.005614 | 40924 | 232 | 5314 | 30 | 1 |
| 11:57070069:C:T | exm910447 | TNKS1BP1 | 0.001706 | 0.005429 | 39870 | 65 | 5313 | 29 | 6.97E-07 |
| 11:70170536:A:G | exm936578 | PPFIA1 | 0.006924 | 0.005447 | 40004 | 282 | 5295 | 29 | 0.2149 |
| 6:46851334:C:T | exm553084 | ADGRF5 (GPR116) | 0.0007885 | 0.00468 | 41334 | 33 | 5317 | 25 | 1.05E-09 |
| 17:8045136:G:A | exm1290985 | PER1 | 0.001434 | 0.003555 | 41282 | 57 | 5325 | 19 | 0.000847 |
| 4:76956246:T:C | exm407053 | CXCL11 | 0.004899 | 0.002995 | 40984 | 204 | 5326 | 16 | 0.05541 |
| 22:19127403:C:T | exm1584742 | ESS2 | 0.0003345 | 0.002808 | 41374 | 13 | 5327 | 15 | 6.33E-08 |
| 19:55143609:G:A | variant.58198 | LILRB1 | NA | 0.002072 | NA | NA | 5297 | 11 | NA |
| 19:6427409:C:T | exm1413868 | SLC25A41 | 4.78E-05 | 0.001878 | 41394 | 2 | 5316 | 10 | 1.95E-08 |
| 3:9988425:T:C | exm289168 | PRRT3 | 0.000339 | 0.001881 | 40834 | 13 | 5306 | 10 | 0.000112 |
| 20:52789469:T:C | exm1551406 | CYP24A1 | 0.000669 | 0.001871 | 41346 | 27 | 5334 | 10 | 0.007187 |
| 19:49351155:A:G | exm1488444 | PLEKHA4 | NA | 0.001706 | NA | NA | 5267 | 9 | NA |
| 18:44221971:C:T | exm1385333 | LOXHD1 | 0.0007646 | 0.001314 | 41336 | 32 | 5321 | 7 | 0.2036 |
| 1:230979494:G:A | exm158330 | C1orf198 | 2.13E-03 | 0.001312 | 41222 | 87 | 5327 | 7 | 0.2587 |
| 13:25479542:C:G | exm1058471 | CENPJ | 0.00399 | 0.001145 | 35870 | 141 | 5232 | 6 | 0.000689 |
| 15:38643456:T:C | exm1147382 | SPRED1 | 0.0004062 | 0.001129 | 41368 | 16 | 5308 | 6 | 0.03235 |
| 17:2227988:T:C | exm2225932 | TSR1 | 0.0004541 | 0.0009409 | 41354 | 19 | 5309 | 5 | 0.1847 |
| 7:63982623:T:C | exm623076 | ZNF680 | NA | 0.0009413 | NA | NA | 5307 | 5 | NA |
