## Supplementary Table 5 for "Rare protein coding variants implicate genes involved in risk of suicide death"

**Table S5**. Prevalence of 30 EMR phenotypes in cases carrying minor allele of rare risk variants.

|  | SNAPC1 14-62244854-C-T | TNKS1BP1 11-57070069-C-T | ESS2 22-19127403-C-T | ADGRF5 6-46851334-C-T | PER1 17-8045136-G-A |
| --- | --- | --- | --- | --- | --- |
| % of total 119 individuals with risk SNPs | 0.28 | 0.24 | 0.13 | 0.21 | 0.16 |
| % of SNP carriers with EMR data | 0.85 | 0.79 | 0.93 | 0.72 | 0.79 |
| Male | 0.89 | 0.91 | 0.79 | 0.83 | 0.73 |
| Female | 0.11 | 0.09 | 0.21 | 0.17 | 0.27 |
| Accidental Trauma | 0.57 | 0.64 | 0.86 | 0.28 | 0.67 |
| Alcohol | 0.21 | 0.18 | 0.36 | 0.28 | 0.20 |
| Anxiety Non Trauma | 0.32 | 0.36 | 0.43 | 0.33 | 0.20 |
| Asthma | 0.32 | 0.36 | 0.43 | 0.33 | 0.20 |
| Bipolar | 0.07 | 0.09 | 0.07 | 0.06 | 0.07 |
| Cardiovascular | 0.46 | 0.45 | 0.21 | 0.39 | 0.40 |
| COPD | 0 | 0.09 | 0.07 | 0.11 | 0 |
| Dementia Neurodegenerative | 0.14 | 0.14 | 0.14 | 0.06 | 0.13 |
| Depression | 0.32 | 0.50 | 0.57 | 0.50 | 0.47 |
| Developmental | 0 | 0.05 | 0.07 | 0.06 | 0 |
| Drug | 0.14 | 0.32 | 0.43 | 0.28 | 0.33 |
| Eating | 0 | 0 | 0 | 0 | 0 |
| Head Injury | 0.11 | 0.14 | 0.07 | 0 | 0.07 |
| Immune Autoimmune | 0.18 | 0.50 | 0.21 | 0.17 | 0.27 |
| Impulse Control Disorders | 0.04 | 0.09 | 0 | 0.06 | 0 |
| Injury By Law Enforcement | 0 | 0 | 0 | 0 | 0 |
| Interpersonal Trauma | 0.25 | 0.50 | 0.36 | 0.22 | 0.53 |
| Migraine | 0.07 | 0.23 | 0.14 | 0 | 0.2 |
| Nutrition Metabolism | 0.32 | 0.41 | 0.29 | 0.33 | 0.47 |
| Obesity | 0.29 | 0.32 | 0.21 | 0.17 | 0.20 |
| Pain | 0.64 | 0.68 | 0.57 | 0.50 | 0.87 |
| PD Cluster A | 0 | 0 | 0 | 0 | 0 |
| PD Cluster B | 0 | 0.09 | 0.14 | 0.06 | 0 |
| PD Cluster C | 0 | 0 | 0 | 0 | 0 |
| Schizophrenia Schizoaffective | 0 | 0.14 | 0.07 | 0 | 0.07 |
| Seizure | 0.04 | 0.05 | 0 | 0 | 0.07 |
| Sleep | 0.36 | 0.32 | 0.43 | 0.17 | 0.20 |
| Somatic | 0.07 | 0.23 | 0 | 0.06 | 0.13 |
| Suicidal Ideation | 0.11 | 0.09 | 0.14 | 0.11 | 0.20 |
| Self Injury | 0.21 | 0.18 | 0.29 | 0.06 | 0.20 |
