## Supplementary figures and images for "Rare protein coding variants implicate genes involved in risk of suicide death"

### Supplementary Figure 1

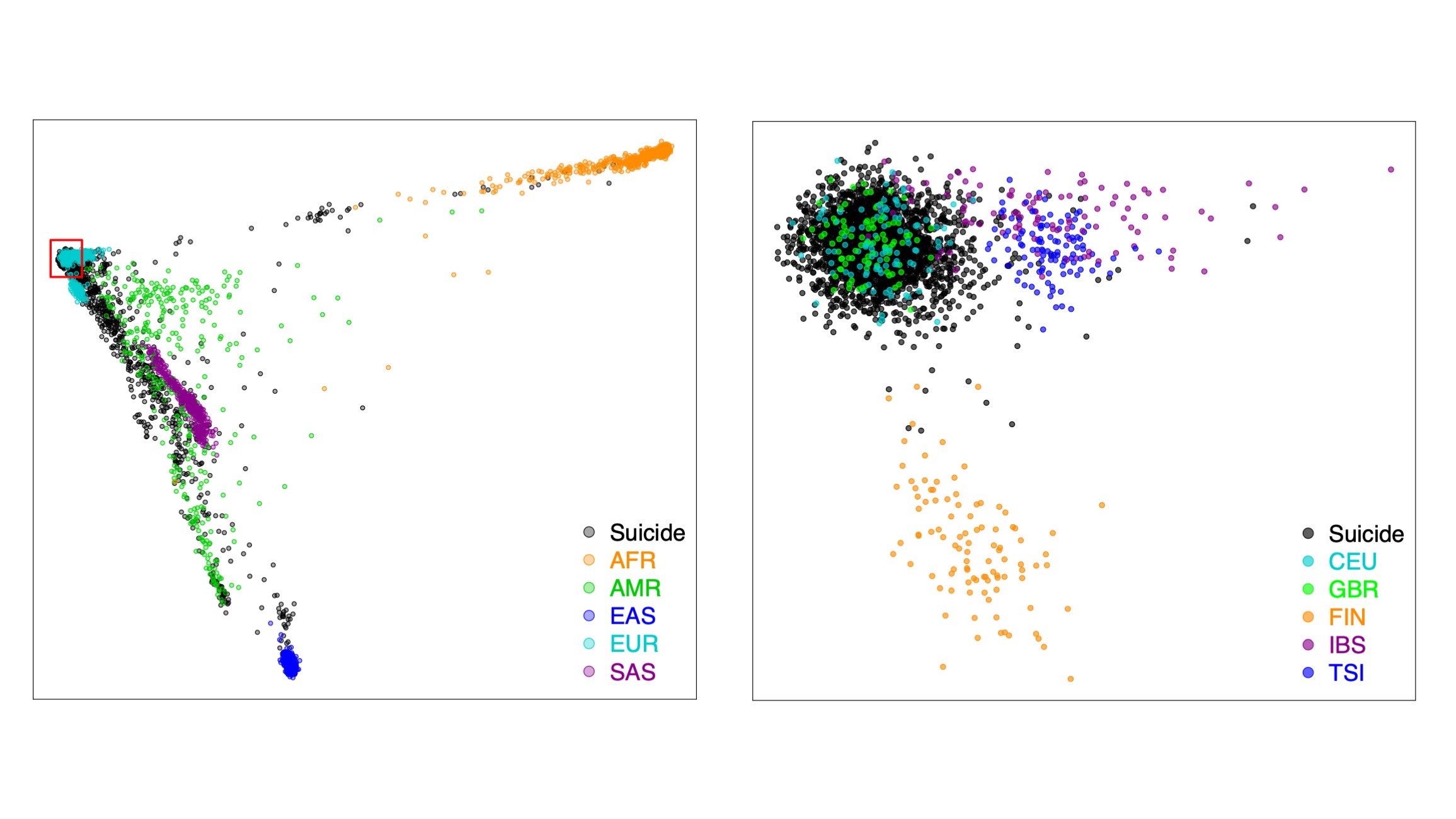
