## Supplementary Material PGC author list for "Rare protein coding variants implicate genes involved in risk of suicide death"

Benjamin M Neale ^1, 2, 3^, Niamh Mullins ^4^, Naomi R Wray ^5, 6^, Rodney J Scott ^7, 8, 9^, Trish Collinson ^7^, Vaughan J Carr ^9, 10^, Bryan J Mowry ^6, 11^, Cynthia Shannon Weickert ^10, 12, 13^, Melissa J Green ^10, 12^, Thomas W Weickert ^9, 10, 12, 14^, Muhammad Ayub ^15^, Brenda J Meyer ^16^, Edmund Sonuga-Barke ^17, 18^, Bernhard T Baune ^19, 20^, Udo Dannlowski ^20^, Claiton H D Bau ^21, 22^, David L Braff ^23^, Barbara Franke ^24, 25^, Christine Lochner ^26^, Damiaan Denys ^27^, Danielle Posthuma ^28, 29^, Eske M Derks ^30^, Roel A Ophoff ^31, 32^, Micha Gawlik ^33^, Andreas Reif ^34^, Carlos N Pato ^35^, Michele T Pato ^35^, Dieter B Wildenauer ^36, 37^, Alessandra Frustaci ^38, 39^, Fabrizio Piras ^40^, Gianfranco Spalletta ^41, 42^, Giuseppe Ducci ^43^, Maria Cristina Cavallini ^44^, Stefano Bonassi ^45^, Stefano Landi ^46^, Alessandro Serretti ^47^, Howard J Edenberg ^48^, John I Nurnberger ^49^, Roy H Perlis ^50, 51^, Israel Liberzon ^52^, Claudio Toma ^12, 53^, Janice M Fullerton ^12, 53^, Peter R Schofield ^12, 53^, Philip B Mitchell ^10^, Andrea L Roberts ^54^, Karestan C Koenen ^54^, Boris Chaumette ^55^, Marie-Odile Krebs ^56, 57, 58^, Douglas F Levinson ^59^, Luis Augusto Rohde ^60, 61^, Claudine Laurent ^62, 63^, Dominique Campion ^64, 65^, Anna Gareeva ^66^, Valentina Escott-Price ^67, 68^, Jordan W Smoller ^2, 69, 70^, Aitana Bigorra ^71^, Amaia Hervas ^71, 72^, Cristina Sánchez-Mora ^73, 74, 75^, Gemma Español Martin ^74^, Isabel Rueda ^76^, Josep Antoni Ramos-Quiroga ^74, 77^, María Jesús Arranz ^78^, María Soler Artigas ^73, 74, 75^, Miquel Casas-Brugué ^79^, Rosa Bosch-Munsó ^74^, Silvina Guijarro ^71^, Vanesa Richarte ^73, 74, 80^, Marta Ribasés ^75, 81, 82^, Diego Albani ^83^, Eduard Vieta ^84^, Luisa Lázaro ^75, 85, 86^, Pino Alonso ^75, 87, 88^, Ingrid Kockum ^89^, Jan Hillert ^90^, Lars Klareskog ^91, 92^, Mikael Landén ^93, 94^, Ming T Tsuang ^95^, Andrew M McIntosh ^96, 97^, Andrew McQuillin ^98^, Arianna Di Florio ^99, 100^, Douglas H R Blackwood ^97^, Gerome Breen ^101, 102^, Nicholas Craddock ^99^, Richard A Bryant ^103^, Joel Gelernter ^104, 105, 106^, Sarah E Medland ^107^, Fernando Goes ^108^, Lynn E DeLisi ^109^, Carol A Mathews ^110^, Dennis L Murphy ^111^, James J Crowley ^112, 113^, Sandra K Loo ^114^, Margaret A Richter ^115^, Irwin D Waldman ^116^, Ian Gizer ^117^

1, Analytic and Translational Genetics Unit, Massachusetts General Hospital, Boston, MA, US

2, Stanley Center for Psychiatric Research, Broad Institute, Cambridge, MA, US

3, Medical and Population Genetics, Broad Institute, Cambridge, MA, US

4, Department of Genetics and Genomic Sciences, Icahn School of Medicine at Mount Sinai, New York, NY, US

5, Institute for Molecular Bioscience, The University of Queensland, Brisbane, QLD, AU

6, Queensland Brain Institute, The University of Queensland, Brisbane, QLD, AU

7, School of Biomedical Sciences, University of Newcastle, Newcastle, NSW, AU

8, Hunter New England Health Service, Newcastle, NSW, AU

9, Schizophrenia Research Institute, Sydney, NSW, AU

10, School of Psychiatry, University of New South Wales, Sydney, NSW, AU

11, Queensland Centre for Mental Health Research, The University of Queensland, Brisbane, QLD, AU

12, Neuroscience Research Australia, Sydney, NSW, AU

13, Department of Neuroscience and Physiology, Upstate Medical University, Syracuse, NY, US

14, Department of Neuroscience and Physiology, State University of New York Upstate Medical University, Syracuse, NY, US

15, Department of Psychiatry, Queen's University, Kingston, ON, CA

16, Psychology, University of Southampton, Southampton, UK

17, Department of Child and Adolescent Psychiatry, King's College London, London, UK

18, Department of Child and Adolescent Psychiatry, Aarhus University,, DK

19, Department of Psychiatry, Melbourne Medical School, University of Melbourne, Melbourne, VIC, AU

20, Department of Psychiatry, University of Münster, Münster, NRW, DE

21, Department of Genetics, Institute of Biosciences, Universidade Federal do Rio Grande do Sul, Porto Alegre, RS, BR

22, ADHD Outpatient Program, Adult Division, Hospital de Clínicas de Porto Alegre, Porto Alegre, RS, BR

23, Department of Psychiatry, University of California, San Diego, CA, US

24, Department of Human Genetics, Donders Institute for Brain, Cognition and Behaviour, Radboud University Medical Center, Nijmegen, GE, NL

25, Department of Psychiatry, Donders Institute for Brain, Cognition and Behaviour, Radboud University Medical Center, Nijmegen, GE, NL

26, MRC Unit on Risk and Resilience in Mental Disorders, Department of Psychiatry, Stellenbosch University, Cape Town, WC, ZA

27, Department of Psychiatry, Academic Medical Centre, Amsterdam, NH, NL

28, Department of Complex Trait Genetics, Center for Neurogenomics and Cognitive Research, Amsterdam Neuroscience, Vrije Universiteit Amsterdam, Amsterdam, NH, NL

29, Department of Clinical Genetics, Amsterdam Neuroscience, Vrije Universiteit Medical Center, Amsterdam, NH, NL

30, Genetics and Computational Biology, QIMR Berghofer Medical Research Institute, Herston, QLD, AU

31, Department of Psychiatry and Biobehavioral Sciences, University of California Los Angeles, Los Angeles, CA, US

32, Center for Neurobehavioral Genetics, Semel Institute for Neuroscience and Human Behavior, University of California Los Angeles, Los Angeles, CA, US

33, Department of Psychiatry, Psychosomatics and Psychotherapy, University of Würzburg, Würzburg, BY, DE

34, Department of Psychiatry, Psychosomatic Medicine and Psychotherapy, University Hospital Frankfurt, Frankfurt am Main, DE

35, Department of Psychiatry and Zilkha Neurogenetics Institute, Keck School of Medicine at University of Southern California, Los Angeles, CA, US

36, School of Psychiatry and Clinical Neurosciences, University of Western Australia, Perth, WA, AU

37, Centre of Clinical Research in Neuropsychiatry, Graylands Hospital, Mt Claremont, WA, AU

38, Eating Disorder Service, St. Ann's Hospital, London, UK

39, Barnet Enfield and Haringey Mental Health NHS Trust, London, UK

40, Department of Clinical and Behavioral Neurology, Neuropsychiatry Laboratory, IRCCS Santa Lucia Foundation, Rome, IT

41, Laboratory of Neuropsychiatry, IRCCS Santa Lucia Foundation, Rome, IT

42, Division of Neuropsychiatry, Menninger Department of Psychiatry and Behavioral Sciences, Baylor College of Medicine, Houston, TX, US

43, Mental Health Department, ASL Roma 1, Rome, IT

44, Department of Neuropsychiatric Sciences, Universita Vita-Salute Ospedale San Raffaele, Milan, IT

45, Clinical and Molecular Epidemiology, IRCCS San Raffaele Pisana, Rome, IT

46, Dipartimento di Biologia, University of Pisa, Pisa, IT

47, Department of Biomedical and NeuroMotor Sciences, University of Bologna, Bologna, IT

48, Biochemistry and Molecular Biology, Indiana University School of Medicine, Indianapolis, IN, US

49, Psychiatry, Indiana University School of Medicine, Indianapolis, IN, US

50, Psychiatry, Harvard Medical School, Boston, MA, US

51, Division of Clinical Research, Massachusetts General Hospital, Boston, MA, US

52, Department of Psychiatry, University of Michigan, Ann Arbor, MI, US

53, School of Medical Sciences, University of New South Wales, Sydney, NSW, AU

54, Harvard University T.H. Chan School of Public Health, Boston, MA, US

55, Department of Neurology and Neurosurgery, McGill University, Montreal, QC, CA

56, Faculté de Médecine Paris Descartes, Université de Paris, Faculté de Médecine Paris Descartes, Paris, France, Paris, FR

57, INSERM, Institut de Psychiatrie et Neurosciences de Paris, Paris, FR

58, GHU Paris Psychiatrie et Neurosciences - Sainte Anne, Service Hospitalo-Universitaire, Paris, FR

59, Psychiatry & Behavioral Sciences, Stanford University, Stanford, CA, US

60, National Institute of Developmental Psychiatry for Children and Adolescents, São Paulo, SP, BR

61, Department of Psychiatry, Program of Developmental Psychiatry, Federal University of Rio Grande do Sul, Hospital de Clinicas de Porto Alegre, Porto Alegre, RS, BR

62, Department of Child and Adolescent Psychiatry, Université Pierre et Marie Curie, Hôpital Pitié-Salpêtrière, Paris, FR

63, Department of Psychiatry, Stanford University, Stanford, CA, US

64, Department of Genetics and CNR-MAJ, Normandy Center for Genomic and Personalized Medicine, Normandie University, Rouen, FR

65, Department of Research, Rouvray Psychiatric Hospital, Sotteville-Lès-Rouen, FR

66, Institute of Biochemistry and Genetics, Ufa Scientific Centre, Russian Academy of Science, Ufa, RF

67, Dementia Research Institute, Cardiff University, Cardiff, UK

68, Division of Psychological Medicine and Clinical Neurosciences, Cardiff University, Cardiff, UK

69, Department of Psychiatry, Massachusetts General Hospital, Boston, MA, US

70, Psychiatric and Neurodevelopmental Genetics Unit (PNGU), Massachusetts General Hospital, Boston, MA, US

71, Child and Adolescent Mental Health Unit, University Hospital Mutua Terrassa, Barcelona, ES

72, Institut Global d'Atenció Integral del Neurodesenvolupament (IGAIN), Barcelona, ES

73, Group of Psychiatry, Mental Health and Addiction, Vall d'Hebron Research Institute, Barcelona, ES

74, Psychiatry Department, Vall d'Hebron University Hospital, Barcelona, ES

75, Biomedical Network Research Centre on Mental Health (CIBERSAM), Madrid, ES

76, Department of Psychiatry, Hospital Sant Joan de Déu, Barcelona, ES

77, Departamento de Psiquiatría y Medicina Legal, Universitat Autònoma de Barcelona, Barcelona, ES

78, University Hospital Mutua Terrassa, Barcelona, ES

79, Servicio de Psiquiatria, Hospital Universitari Vall d' Hebron, Universitat Autònoma de Barcelona, Barcelona, ES

80, Biomedical Network Research Centre on Mental Health (CIBERSAM), Barcelona, ES

81, Department of Psychiatry, Vall d'Hebron University Hospital, Barcelona, ES

82, Psychiatric Genetics Unit, Vall d’Hebron Research Institute, Universitat Autònoma de Barcelona, Barcelona, ES

83, Neuroscience Department, Istituto di Ricerche Farmacologiche Mario Negri IRCSS, Milan, IT

84, Bipolar and Depressive Disorders Unit, Hospital Clinic, Institute of Neurosciences, University of Barcelona, Barcelona, ES

85, Department of Child and Adolescent Psychiatry and Psychology, Institute of Clinical Neurosciences, Barcelona, ES

86, IDIBAPS, Barcelona, ES

87, Psychiatry Department, Bellvitge University Hospital, Bellvitge Biomedical Research Institute (IDIBELL), Bacelona, ES

88, Department of Clinical Sciences, University of Barcelona, Barcelona, ES

89, Neuroimmunology Unit, Department of Clinical Neuroscience, Karolinska Institutet, Stockholm, SE

90, Division of Neuroscience, Department of Clinical Neuroscience, Karolinska Institutet, Stockholm, SE

91, Institute of Environmental Medicine, Karolinska Institutet, Stockholm, SE

92, Centre of Occupational and Environmental Medicine, Stockholm County Council, Stockholm, SE

93, Department of Medical Epidemiology and Biostatistics, Karolinska Institutet, Stockholm, SE

94, Institute of Neuroscience and Physiology, University of Gothenburg, Gothenburg, SE

95, University of California San Diego, La Jolla, CA, US

96, Centre for Cognitive Ageing and Cognitive Epidemiology, University of Edinburgh, Edinburgh, GB

97, Division of Psychiatry, University of Edinburgh, Edinburgh, GB

98, Division of Psychiatry, University College London, London, GB

99, Medical Research Council Centre for Neuropsychiatric Genetics and Genomics, Division of Psychological Medicine and Clinical Neurosciences, Cardiff University, Cardiff, GB

100, Department of Psychiatry, University of North Carolina at Chapel Hill, Chapel Hill, NC, US

101, MRC Social, Genetic and Developmental Psychiatry Centre, King's College London, London, GB

102, NIHR BRC for Mental Health, King's College London, London, GB

103, University of New South Wales, Sydney, NSW, AU

104, Division of Human Genetics, Department of Psychiatry, Yale University School of Medicine, New Haven, CT, US

105, VA CT Healthcare Center, West Haven, CT, US

106, Departments of Genetics and Neuroscience, Yale University School of Medicine, New Haven, CT, US

107, Genetic Epidemiology, QIMR Berghofer Medical Research Institute, Brisbane, QLD, AU

108, Department of Psychiatry and Behavioral Sciences, Johns Hopkins University School of Medicine, Baltimore, MD, US

109, Department of Psychiatry, Cambridge Health Alliance, Harvard Medical School, Cambridge, MA, US

110, Department of Psychiatry, University of Florida, Gainesville, FL, US

111, Laboratory of Clinical Science, National Institute of Mental Health Intramural Research Program, National Institutes of Health, Bethesda, MD, US

112, Center for Psychiatric Genomics, Chapel Hill, NC, US

113, Department of Genetics, University of North Carolina, Chapel Hill, NC, US

114, Semel Institute for Neuroscience and Human Behavio, University of California, Los Angeles, Los Angeles, CA, US

115, Frederick W. Thompson Anxiety Disorders Centre, Sunnybrook Health Sciences Centre, University of Toronto, Toronto, ON, CA

116, Department of Psychology, Emory University, Atlanta, GA, US

117, Department of Psychological Sciences, University of Missouri, Columbia, MO, US
